# Evaluation of direct strain field prediction in bone with data-driven image mechanics (D^2^IM-Strain)

**DOI:** 10.64898/2026.03.31.715417

**Authors:** Jon Valijonov, Peter Soar, James Le Houx, Gianluca Tozzi

## Abstract

Digital volume correlation (DVC) has become the benchmark experimental technique for full-field strain measurement in bone mechanics. In our previous work we developed a novel data-driven image mechanics (D^2^IM) approach that learns from DVC data and predicts displacement fields directly from undeformed X-ray computed tomography (XCT) images, deriving strain fields from such predictions. However, strain fields derived through numerical differentiation of displacement fields amplify high-frequency noise, and regularization techniques compromise spatial resolution while incurring substantial computational costs. Here we propose the upgrade D^2^IM-Strain to predict strain fields directly from XCT images of bone. Two prediction strategies were compared: displacement-derived strain and direct strain prediction. The direct strain prediction model significantly improved accuracy particularly for strain magnitudes below 10000με, taken as a representative threshold value for bone tissue yielding in compression. In addition, the direct approach reduced false-positive high-strain classifications by 75%. By eliminating numerical differentiation, the approach reduces noise amplification while maintaining computational efficiency. These findings represent a critical step toward developing robust data-driven volume correlation methods for hierarchical materials.

## 1. Introduction

Hard tissues exhibit a complex hierarchical architecture, characterised by significant heterogeneity in their mechanical properties across multiple scales. This complexity poses challenges for their mechanical evaluation due to the need to account for both scale-dependent behaviour and structural heterogeneity [1–4]. In response, mechanical characterisation methodologies have evolved from direct experimental approaches such as macro-to-nano mechanical testing to advanced techniques that incorporate multi-scale observations within a single analysis framework. Digital volume correlation (DVC) has now been established as a benchmark technique in the mechanical assessment of hard tissues [5, 6]. By leveraging three-dimensional (3D) images mainly obtained through high-resolution X-ray computed tomography (XCT) at multiple steps of deformation *in situ*, DVC enables full-field measurements of displacement and strain within the material [7, 8].

DVC methodologies can be broadly classified into local and global approaches [9], both of which are widely adopted through commercial software, freeware, or custom solutions. Both approaches have been extensively validated for bone mechanics applications [10] and face limitations, particularly in the derivation of strain fields [11, 12]. The calculation of strain requires numerical differentiation of the displacement field, a process that amplifies high-frequency noise and introduces errors [13–15]. To mitigate these issues, existing techniques impose regularization either by smoothing or by making assumptions about the mechanical properties of the tissue being imaged [5]. In fact, even advanced regularization techniques, such as diffeomorphic smoothing, hyperelastic warping, or finite element-based methods remain computationally expensive and compromise the resolution of strain measurements. The selection of appropriate regularisation parameters represents a fundamental trade-off between noise suppression and the preservation of spatial features, with parameter choices often requiring empirical tuning for each experimental configuration [16]. To overcome such issues, a 3D direct deformation estimation approach was proposed to estimate deformation gradient fields directly from image volumes without explicit consideration of displacement fields [17].

However, these algorithms require an initial guess that can cause the optimisation to return an incorrect solution or fail to converge, with consequent implications for both spatial and temporal resolution. These challenges underscore the need for innovative approaches that can enhance strain field accuracy without sacrificing resolution or computational efficiency.

The advent of deep learning techniques has opened new possibilities for full-field displacement and strain determination in material characterisation. Convolutional neural networks (CNNs) offer the ability to learn complex patterns from data, making them attractive for predicting displacement and strain fields directly from image inputs [18]. As inherently data-driven models, however, CNNs may encode specimen-specific morphological or imaging characteristics, raising questions regarding transferability to unseen domains. Nevertheless, strategies such as data augmentation and normalisation can substantially improve cross-domain robustness [19–21]. In addition to these considerations, the application of CNNs for strain field computation introduces further challenges. When strain fields are derived through numerical differentiation of CNN-predicted displacement fields, high-frequency noise is often magnified. While Gaussian filters are commonly applied to smooth displacement fields prior to strain calculation [22], this approach diminishes the advantage of CNNs in capturing fine spatial details and high-frequency deformation features.

To address this limitation, frameworks such as StrainNet have been developed to enable direct strain prediction from input images, bypassing displacement field computation entirely [23]. By eliminating the need for numerical differentiation, StrainNet minimises noise amplification, enabling high spatial resolution in strain predictions without requiring post-processing filters. Additionally, the deep learning architecture inherently accounts for and removes the influence of rigid body motions and translations during strain computation. Despite these advantages, StrainNet’s application has been limited to two-dimensional (2D) strain prediction from speckled images, primarily within the scope of digital image correlation (DIC). Its architecture and training strategy are tailored to 2D surface deformation, and this limitation motivates the need for approaches that operate on volumetric images and enable direct strain prediction beyond the constraints of surface-based DIC.

Building on this foundation, we recently introduced a novel CNN-based framework known as data-driven image mechanics (D^2^IM), designed to predict displacement fields directly from high-resolution XCT images of hard tissues [24]. The D^2^IM framework was validated using XCT datasets of vertebral bone, including both intact and artificially lesioned specimens. To enhance dataset size, 2D cross-sectional slices from the XCT tomograms were used, allowing the model to improve accuracy for those imaging and loading conditions. The D^2^IM framework demonstrated good predictive performance for displacement fields in the primary loading direction of the vertebrae, even when applied to lower-resolution clinical images. However, as the D^2^IM approach derives strain fields by differentiating the predicted displacement fields, the resulting strain predictions inherit displacement-related errors. These errors are primarily reflected as underestimations of higher strain magnitudes and overestimations of lower strain magnitudes.

To overcome such shortcomings, we propose a novel output strategy within the D^2^IM framework to enable direct strain prediction (D^2^IM-Strain). This strategy bypasses the numerical differentiation of displacement fields, aiming to improve the accuracy of strain predictions. Our findings demonstrate that direct strain prediction within the D^2^IM-Strain model outperforms strain fields derived from predicted displacement, resulting in a reduced relative error distribution. This improvement is particularly pronounced for strain magnitudes below the yield threshold of bone in compression (i.e. 10000με) [25], a range that encompasses most of the bone tissue under both physiological loading conditions. This study represents a critical step toward the development of data-driven volume correlation within the broader D^2^IM framework. By leveraging deep learning to achieve high-resolution, noise-resistant strain predictions, this approach has the potential to advance the mechanical characterisation of hierarchical materials such as bone.

## 2. Methods

The tomography dataset, DVC calculation, and model architecture have been described in detail in our previous work [24], where the network architecture was intentionally kept as close as possible to the original D^2^IM framework to enable a direct and fair comparison between displacement and strain prediction tasks (Fig. 1).

**Figure 1:**
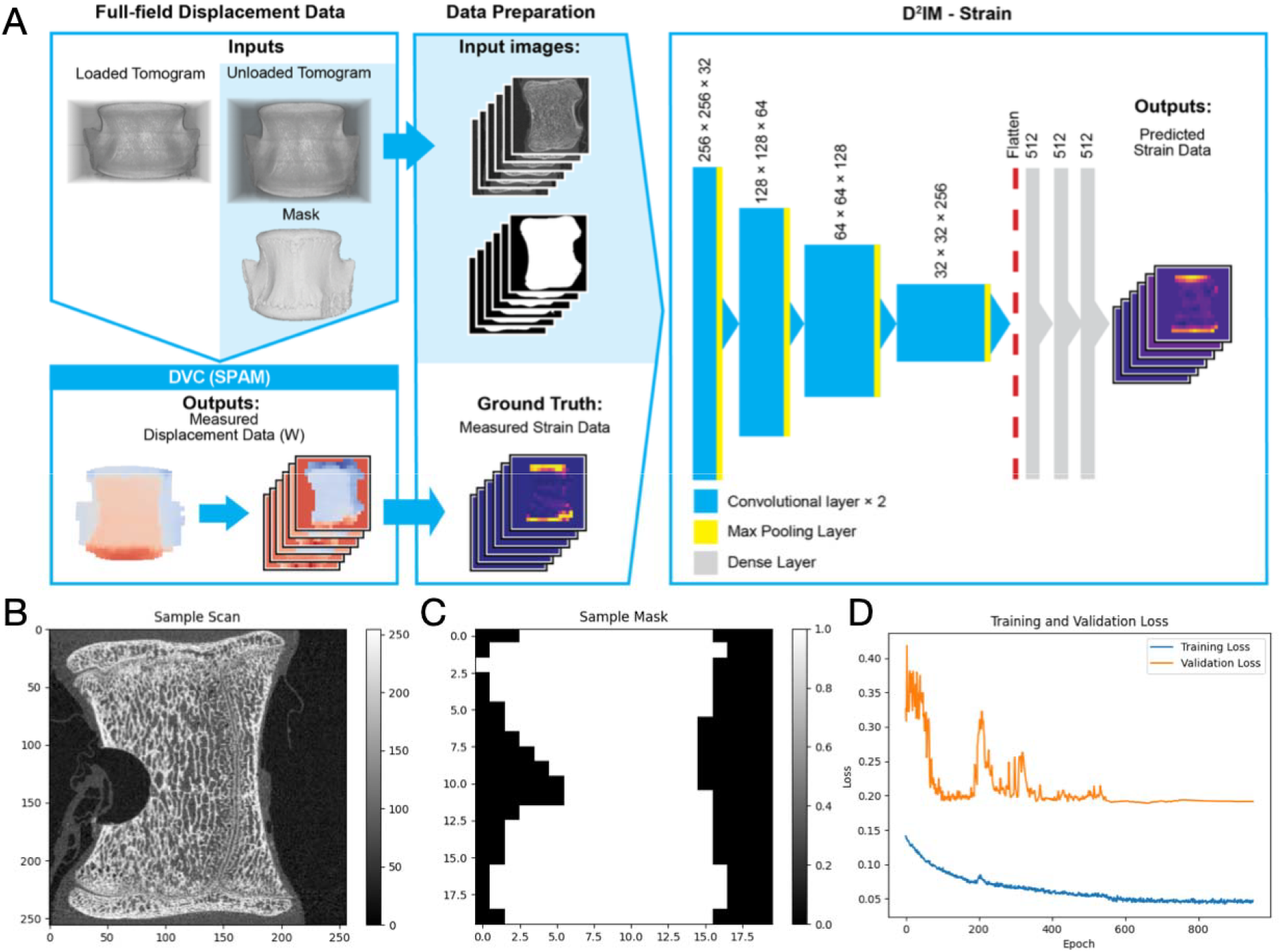
(A) Workflow depicting the strategy for direct prediction of strain fields straight from D^2^IM-Strain image input. Workflow for direct strain field prediction within the D^2^IM framework. (Left) Data preparation and ground truth generation: DVC is applied to loaded and unloaded XCT scans to calculate displacement fields, which are numerically differentiated to generate ground-truth strain labels. (Right) The D^2^IM CNN architecture. The model inputs only the unloaded tomogram and binary mask to directly predict strain. The architecture (4 convolutional blocks with max-pooling, followed by 3 dense layers with ReLU activation) was kept identical to the original displacement-based D^2^IM to ensure a direct, fair comparison between the two prediction strategies. (B) Sample greyscale scan with lesion. (C) Mask generated from scan in (B). (D) Training process of the model, illustrating performance across epochs.

Briefly, we used a publicly available dataset [26] consisting of 10 porcine vertebrae (5 lesioned and 5 intact), each scanned using *in situ* XCT under two loading conditions (unloaded and loaded) with an isotropic voxel size of 39 μm [27, 28]. For the lesioned group, artificial defects (approx. 5 mm diameter and 5 mm depth) were created in the central region of the vertebral body using a dental drill to simulate focal bone loss, representing simplified models of metastatic or osteoporotic lesions. DVC displacement and strain fields were measured using the open-source SPAM Python library [29], which implements a local DVC approach of sufficient accuracy for such applications [30]. A binary mask was generated from the unloaded scan for each image using ImageJ [31], automatically defining the binary threshold based on a slice in the centre of the structure, applying a dilation of 8 voxels, and filling in any remaining internal holes. For DVC, a window size of 50 voxels was selected and deemed sufficiently large to capture multiple trabeculae while maintaining spatial resolution adequate to resolve local strain variations.

The tomograms were sliced into 420 two-dimensional images extracted perpendicular to the loading axis. Slices lacking sufficient structural detail, such as those containing predominantly background or cortical shell with minimal trabecular structure, were manually removed through visual inspection, leaving a final dataset of 251 images. This slicing strategy serves a dual purpose: it substantially increases the effective training dataset size without requiring additional experimental acquisitions, and it allows the model to learn from diverse anatomical cross-sections spanning the full vertebral height. All input images were resized to 256 × 256 pixels using bilinear interpolation to standardise input dimensions for the CNN, while the binary mask and predicted strain field were represented on a coarser 20 × 20 grid corresponding to the DVC node spacing. This multi-resolution approach balances the need for high-resolution input features (captured in the 256 × 256 image) with the coarser spatial resolution inherent to DVC measurements (reflected in the 20 × 20 output grid). Grayscale values were normalised to the range 0–1 by dividing by the maximum possible intensity value before being input into the CNN, ensuring consistent input scaling across all specimens and preventing numerical instabilities during training. The ground-truth strain fields used for supervised training were obtained from SPAM-computed displacement fields as described above and served as target labels for the CNN during training. Similarly to [24], we focused on both quantitative and qualitative comparisons of the normal strains in z (*ε*_zz_), which is the axis of largest displacement due to the experimental setup. Standardisation proved most effective, converting strain values to zero mean and unit variance within the region of interest defined by the mask. Values outside the mask were set to zero, ensuring that only physically relevant regions contributed to learning and preventing the model from learning spurious patterns in background regions.

The D^2^IM-Strain architecture is a feed-forward CNN that predicts strain fields directly from the unloaded tomogram and binary mask. The architecture consists of four convolutional blocks followed by three dense layers, identical to the original displacement-prediction D^2^IM model [24] (inspired by VGGNet [32]) to enable direct comparison. Each convolutional block comprises a 2D convolutional layer with 3×3 kernels, ReLU activation, and 2×2 max-pooling for spatial downsampling. The number of filters doubles with each successive block (32, 64, 128, 256), allowing the network to learn increasingly abstract and hierarchical features. The output of the fourth convolutional block is flattened into a one-dimensional feature vector, which is then passed through three fully connected (dense) layers of 512 channels, again using ReLU activation. The final layer outputs corresponding to the flattened 20 × 20 strain field. The binary mask is concatenated with the grayscale image as a second input channel, allowing the network to explicitly learn the spatial extent of the bone region and focus predictions accordingly.

The dataset was shuffled before training to disrupt the spatial correlation between sequential slices and reduce the risk of the model overfitting to the anatomical progression within individual vertebrae. The data were split into training, validation, and test sets using a 60/20/20 ratio (approximately 150 training, 50 validation, 51 test images), ensuring that slices from the same vertebra were not split across training and test sets to avoid data leakage. Initial model performance yielded R^2^=0.50 on the test set. L2 regularisation with a penalty coefficient of 1 × 10^−6^ was then introduced to reduce overfitting and a dropout rate of 0.2 was applied to the dense layers to encourage robust feature learning. These regularisation strategies yielded R^2^=0.55, indicating improved generalization to unseen data. The model was trained using a staged learning rate schedule designed to balance rapid initial convergence with fine-tuning in later epochs. An initial learning rate of 1 × 10^−3^ was applied for the first 600 epochs, allowing the model to quickly navigate the loss landscape. This was followed by a reduced learning rate of 1 × 10^−4^ for 200 epochs to refine the solution, and a final phase of 200 epochs with a learning rate of 1 × 10^−5^ for fine-grained optimization. The model was trained using the Mean Absolute Error (MAE) loss function, which computes the average absolute difference between predicted and ground-truth strain values across all output nodes. MAE was compared to Mean Squared Error (MSE), to assess generalisation of strain distribution. The Adam optimiser was then employed for gradient-based weight updates due to its adaptive learning rate properties and robust performance across diverse architectures [33]. Model training was performed on an NVIDIA RTX A6000 GPU and required approximately 11 minutes and 36 seconds in total for the full 1,000-epoch training cycle. Comparison with the displacement-derived strain approach from our prior work [24] was performed using identical test specimens and evaluation metrics to ensure a fair and direct assessment of the two prediction strategies. The distribution of relative strain errors was compared between the direct strain prediction model and the displacement-derived D^2^IM approach using the Mann-Whitney U test (α = 0.01, scipy.stats, Python).

## 3. Results and Discussion

This study explores strategies to directly predict strain fields using D^2^IM, thereby bypassing numerical differentiation and eliminating the associated propagation of error. As shown in Figure 2A, the direct strain model improves prediction accuracy compared to the displacement-derived strains obtained from the original D^2^IM framework [24]. The direct strain prediction model achieved an R^2^ of 0.55, representing an improvement over the displacement-derived approach. This improvement is particularly evident in the pre-yield region (strain magnitudes below 10000με, corresponding to the tissue yielding value for bone in compression [25]), as shown in Fig. 2B. By bypassing numerical differentiation, the new model significantly reduces strain error in the elastic region (p < 0.01), where traditional displacement-derived methods typically introduce the most noise, but not above yield (p > 0.01). The reduced scatter in the low-strain regime reflects the model’s ability to directly learn the mapping from microstructural features to local strain without the intermediate step of displacement field computation and subsequent differentiation, each of which introduces cumulative errors. While both models correctly identify most low-strain regions (<10000με), the direct strain model substantially reduces the number of false positive predictions (Figure 2C). False positives decreased by almost 75% (from 304 to 80), indicating a stronger ability to correctly distinguish low-strain regions and avoid overestimating high-strain values.

**Figure 2:**
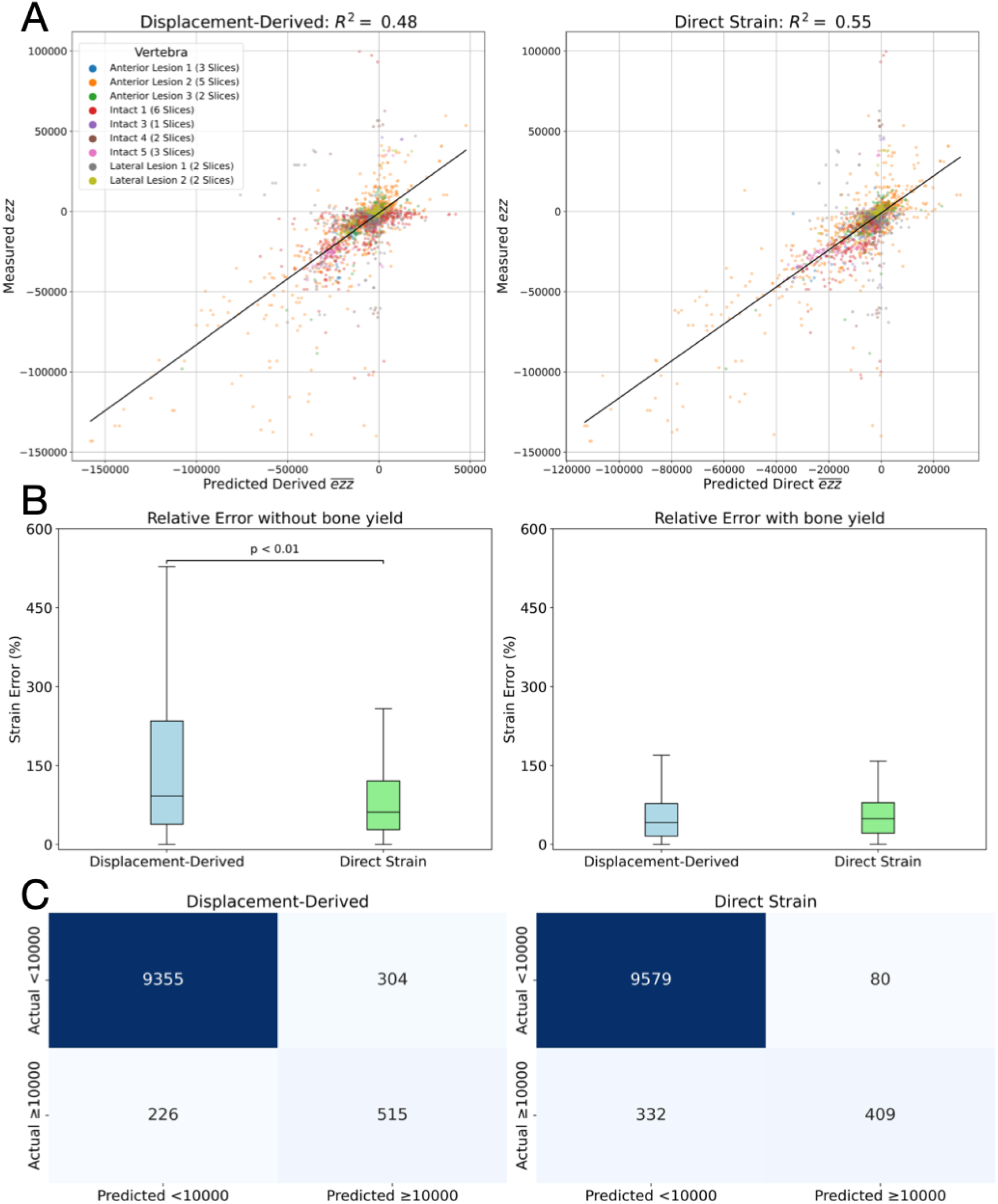
(A) Correlation analysis between all measured and predicted normal strains (displacement derived in [24] vs new model) in the loading direction z (*ε* _zz_) for the test dataset, including a line of best fit and the calculated correlation coefficient R^2^. Points have been coloured according to the vertebra they belong to. (B) Box and whisker chart with relative error distribution as a percentage of the measured *ε*_zz_. Boxes show the interquartile range and whiskers the maximum and minimum observed errors (excluding outliers) for a) only data points <10000με and b) all data points >10000με. (C) Actual vs predicted strain data points for displacement derived in [24] (left) compared to direct prediction from new model (right).

This reduction in false positives is particularly important for applications in bone mechanics, where overestimation of high-strain regions could lead to incorrect identification of tissue at risk of damage or failure. Although the displacement-derived model identifies a slightly larger number of true high-strain cases (515 vs 409; 25% higher), the direct strain model demonstrates a better trade-off in predictions, favouring correct classification of low-strain regions while still capturing a substantial portion of high-strain events.

To understand the factors contributing to improved generalisation, we systematically evaluated the effects of different loss functions and regularisation strategies. As illustrated in Table 1, the model trained using MSE with a dropout rate of 0.2 and L2 regularisation of 1 × 10^−6^ achieved the highest training performance (R^2^=0.97). However, the validation and test scores were substantially lower (R^2^=0.60 and 0.41, respectively), indicating significant overfitting. The large gap between training and test performance suggests that MSE’s quadratic penalty on errors causes the model to overfit to outliers and noise in the training data, learning specimen-specific patterns that do not generalise to unseen vertebrae. A similar trend was observed when using the Huber loss function, where the training performance remained high (R^2^=0.91), but the test performance improved only marginally (R^2^=0.43). The Huber loss, which combines quadratic and linear penalties, provides some robustness to outliers but still exhibited substantial overfitting in this application [34].

**Table 1.** (E) Comparison of model R^2^ scores for training, validation, and test datasets using different loss functions (MSE, Huber, MAE) and regularization settings (dropout and L2).

| Loss | Dropout | L2 | $R^2$ Train | $R^2$ Validation | $R^2$ Test |
| --- | --- | --- | --- | --- | --- |
| MSE | 0.2 | $1e-06$ | 0.97 | 0.60 | 0.41 |
| Huber | 0.2 | $1e-06$ | 0.91 | 0.58 | 0.43 |
| MAE | 0.2 | $1e-06$ | 0.74 | 0.70 | 0.55 |
| MAE | 0.5 | $1e-05$ | 0.80 | 0.60 | 0.46 |

Using MAE with moderate regularisation (dropout = 0.2, L2 = 1 × 10^−6^) produced the best overall performance. In this configuration, the model achieved R^2^ values of 0.74, 0.70, and 0.55 for the training, validation, and test datasets, respectively. The relatively small gap between training and test performance indicates good generalisation, suggesting that this level of regularisation provides an effective balance between model flexibility and overfitting control. To further investigate the role of regularisation, we increased both the dropout rate (0.5) and L2 penalty (1 × 10^−5^) while using MAE. This led to a reduction in performance, with the test R^2^ decreasing from 0.55 to 0.46, indicating that excessive regularisation constrained the model and resulted in underfitting.

Compared with MSE and Huber losses, which achieved higher training R^2^ values but lower test performance, the MAE loss function consistently improved generalisation. This is likely due to its reduced sensitivity to the high-magnitude and localised maximum strain values present in yielding regions, as well as to outliers in the strain data arising from DVC measurement errors in bone strain distributions [11, 13].

Figure 3 presents representative examples comparing the measured strain fields obtained from DVC with the predictions generated by the displacement-derived D^2^IM model and the proposed D^2^IM-Strain model. For each example, the input tomography slice is shown alongside the corresponding measured strain field, predicted strain field, and the relative prediction error. In the case of the intact vertebra (Fig. 3A), both models capture the overall spatial distribution of the normal strain component *ε*_zz_, along the loading direction. Regions of elevated strain near the superior and inferior endplates are reproduced by both approaches, reflecting the concentration of load transfer through these anatomical structures. However, the D^2^IM-Strain model provides a closer qualitative agreement with the measured strain field, particularly in the localisation and magnitude of the peak strain regions. The strain distribution exhibits characteristic patterns associated with vertebral architecture, with higher strains concentrated in regions of lower bone volume fraction and more complex trabecular connectivity [35]. The corresponding relative error maps indicate that prediction errors are generally distributed across the interior of the vertebra, with lower errors observed in regions of higher strain magnitude. This pattern suggests that the model learns more effectively from high-strain examples, which provide stronger training signals, while low-strain regions with smaller absolute values contribute less to the loss function and therefore receive less emphasis during training. A similar trend is observed for the lesioned vertebra (Fig. 3B). The measured strain field exhibits a pronounced concentration at the lesion site, reflecting the altered load distribution caused by the structural defect. The lesion acts as a stress concentrator, redistributing load to the surrounding intact trabecular network and creating steep strain gradients at the lesion boundary. While the displacement-derived D^2^IM model captures the general pattern of strain localisation, the D^2^IM-Strain model more accurately reproduces the spatial extent and intensity of the high-strain region. This improvement is reflected in the predicted strain maps, where the D^2^IM-Strain results show a stronger correspondence with the measured strain distribution, including better resolution of the strain gradient surrounding the lesion. The error maps further demonstrate that the largest prediction deviations occur near areas of sharp strain gradients, particularly around the lesion boundary. This is expected, as steep gradients represent challenging features for any predictive model, and the spatial resolution of the 20 × 20 output grid may be insufficient to fully resolve the finest details of strain localisation. These qualitative comparisons indicate that the direct strain prediction approach can capture the dominant spatial features of the experimentally measured strain fields. In particular, the D^2^IM-Strain model demonstrates improved representation of localised strain concentrations, suggesting that directly learning strain fields from tomographic data provides a more robust mapping between bone microstructure and mechanical response than the intermediate displacement-based approach.

**Figure 3:**
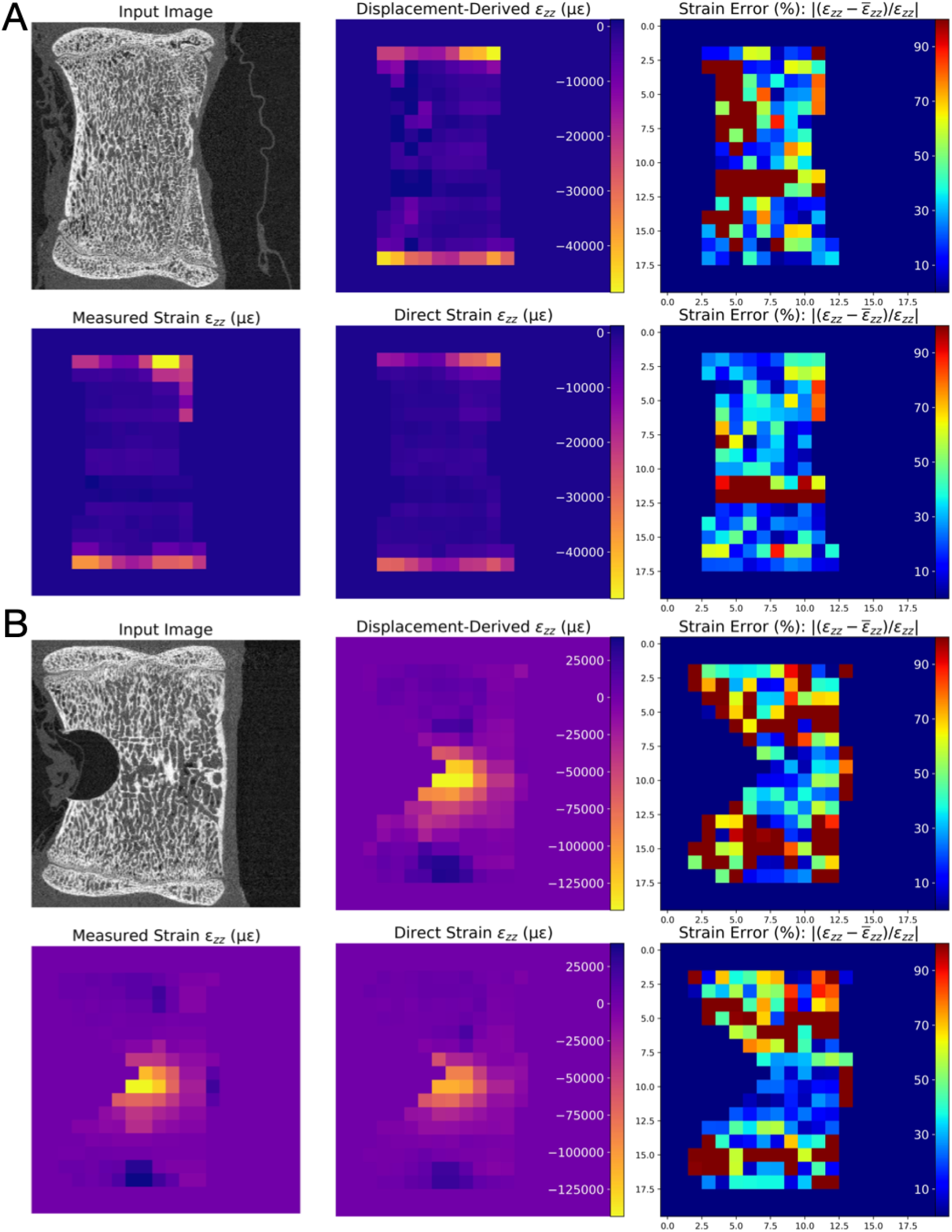
(A) Results for a left-right sliced tomography with no lesion. Input tomography slice, predicted and relative error of *ε* _zz_ from: D^2^IM (top row mid-right) [24], D^2^IM-Strain (bottom row mid-right). (B) Results for a left-right sliced tomography with lesion. Input tomography slice, predicted and relative error of *ε*_zz_ from: D^2^IM (top row mid-right) [24], D^2^IM-Strain (bottom row mid-right).

Figure 4 presents the distribution of relative prediction error across four representative tomography slices from vertebrae, including the two illustrated in Figure 3. The three distinct categories of DVC window behaviour are highlighted using coloured boundary markers to contextualise model performance in relation to bone yielding. In the top row, regions marked in black indicate areas where both the measured DVC strain and the predicted strain are below 10000με. These regions represent correctly identified low-strain zones, demonstrating that the model reliably captures areas experiencing relatively small deformation. The low relative errors observed in these regions suggest that the model maintains stable predictions within structurally intact portions of the vertebra, where strain magnitudes are typically within the elastic regime and trabecular architecture remains mostly undamaged. The middle row, highlighted in white, corresponds to regions where the measured strain remains below 10000με while the predicted strain exceeds this threshold. These areas represent false positive predictions, where the model overestimates the strain magnitude. Such discrepancies are typically located near regions of higher strain gradients, where the transition from low to high strain occurs over a short spatial distance. However, these occurrences are relatively limited compared with the correctly classified regions shown in the other rows. This indicates that the model rarely misclassifies low-strain regions as high-strain regions, demonstrating a strong ability to avoid overestimating strain magnitudes across the vertebral structure. The sparsity of false positives is a key advantage of the direct strain prediction approach, as it reduces the risk of incorrectly identifying tissue as being at risk of yielding or damage.

**Figure 4:**
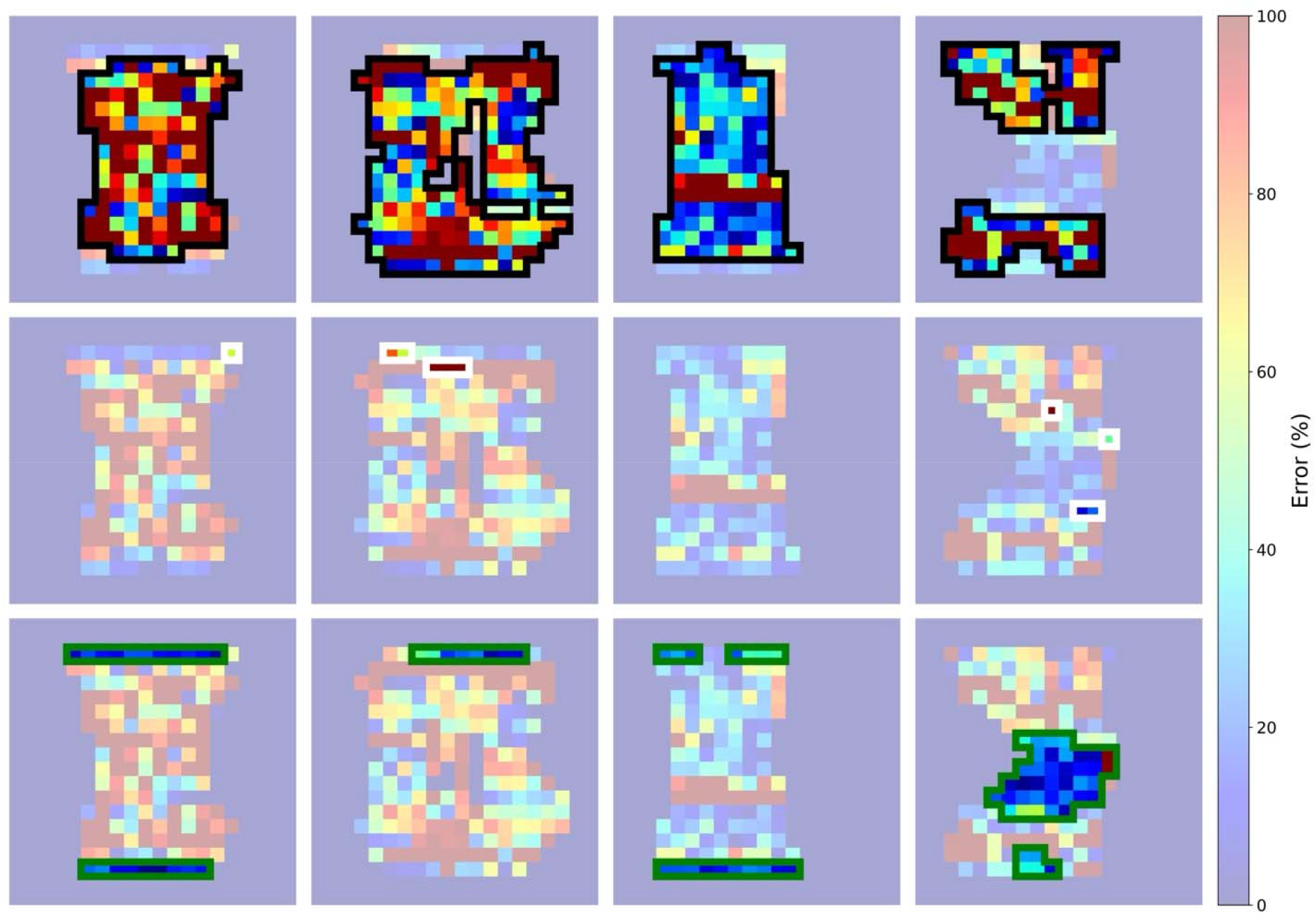
Relative error of strain predictions for the different cases reported in Figure 3 (left-right). Images have been marked with windows highlighting regions of interest regarding bone yielding behaviour. Top: marked in black is measured strain in DVC windows lower than 10000με and predicted strain also lower than 10000με. Middle: marked in white is measured strain in DVC windows lower than 10000με but predicted strain higher than 10000με. Bottom: marked in green is measured strain in DVC windows higher than 10000με and predicted strain also higher than 10000με.

In the bottom row, regions highlighted in green correspond to locations where both the measured and predicted strain values exceeded 10000με. These regions represent correctly identified high-strain zones that may be associated with the onset of bone yielding. The presence of these correctly predicted high-strain regions indicates that the model can capture areas of concentrated mechanical loading, particularly near structural features such as endplates or lesion boundaries. The spatial distribution of these high-strain regions is consistent with our previous studies [24, 35]. The highlighted regions demonstrate that the model can reproduce the dominant mechanical behaviour of the vertebral structure, correctly identifying both low-strain and high-strain zones in most cases. The remaining discrepancies are primarily associated with transitional regions where strain gradients are high, reflecting the inherent challenges of predicting localised strain variations from tomographic data, which is strictly related to image resolution, noise/artifacts and spatial resolution of the measurement [36, 37]. The improved performance of the direct strain prediction approach can be attributed to several factors. First, by eliminating the numerical differentiation step, the model avoids the amplification of high-frequency noise inherent to derivative calculations. In traditional DVC workflows, displacement fields are first computed through image correlation, and strains are subsequently derived by taking spatial derivatives. This two-step process compounds errors: noise in the displacement field is amplified during differentiation, and the resulting strain field requires additional smoothing or regularization [13, 16]. The direct strain prediction approach circumvents this cascade of errors by learning the mapping from image features to strain in a single end-to-end framework. Second, the CNN architecture inherently incorporates spatial context through its hierarchical feature extraction, allowing the model to learn relationships between local microstructure and resulting strain patterns without explicit mechanical assumptions. The convolutional layers capture multi-scale features, from fine trabecular details to larger-scale architectural patterns, and the fully connected layers integrate these features to produce spatially coherent strain predictions. This data-driven approach is particularly advantageous for heterogeneous materials like bone, where constitutive relationships are complex and vary spatially [3, 4].

Despite the promising results, some limitations must be acknowledged. First, the current implementation operates on 2D cross-sectional slices rather than full 3D volumes, sacrificing through-thickness spatial context and the ability to predict all six strain tensor components. Extension to fully volumetric 3D predictions will require architectural modifications, such as replacing 2D convolutional layers with 3D convolutions [38], and will substantially increase computational and memory requirements. Recent work on volumetric optical flow networks, such as VolRAFT [39], provides a promising architectural foundation for this extension. Second, the model’s performance for high-strain regions (>10000με) is limited by the underrepresentation of yielding voxels in the training data. Most bone tissue under typical loading conditions remains in the elastic regime, resulting in a training distribution heavily skewed toward low and moderate strain values. This class imbalance can be addressed through targeted data augmentation strategies, such as oversampling high-strain examples or generating synthetic high-strain data through image generators or finite element simulations, though care must be taken to ensure that synthetic data accurately represents real mechanical behaviour. Third, the present study focuses exclusively on the normal strain component εzz in the primary loading direction, as this component represents the dominant deformation mode in uniaxial compression. However, complete mechanical characterisation requires extension to multi-component strain prediction. This highlights an important trade-off between displacement- and strain-based modelling approaches. Predicting the full displacement field provides greater flexibility, as all strain components can subsequently be derived from the three displacement components. In contrast, direct strain prediction is inherently task-specific: predicting additional strain components would require either multiple specialised models or a single model with substantially expanded output layers. Therefore, while direct strain prediction may offer advantages when a specific strain metric is of primary interest, displacement-based approaches retain broader applicability for deriving arbitrary strain components and enabling more general mechanical analyses. Fourth, the current model was trained and tested on porcine vertebrae under uniaxial compression, and its performance on other anatomical sites, species, loading modes, or pathological conditions is unknown. Systematic validation across diverse datasets will be necessary to assess cross-domain transferability, where transfer learning approaches based on limited data [40] offer a promising avenue for improving generalisability while minimising the need for extensive new experimental acquisitions.

Fifth, the ground-truth strain fields used for training are themselves derived from DVC, meaning that the model inherits any systematic biases or limitations present in such measurements. DVC is subject to well-characterised sources of error, including image noise, limited spatial resolution, and regularisation artifacts [11–13, 16]. Validation against independent ground-truth sources, such as synthetic datasets with known analytical solutions, would provide additional confidence in model predictions and help distinguish between model errors and ground-truth errors. Finally, the current study is limited to mineralised tissue with relatively high X-ray contrast and consistent grayscale appearance. Extension to soft tissues, or biomaterials with lower contrast may require architectural modifications to ensure robust feature extraction.

## 4. Conclusion

This study aimed as predicting strain fields in vertebral bone directly from a single and undeformed XCT image using the D^2^IM-Strain framework. To address the limited availability of experimental training data in experimental mechanics, a dataset augmentation strategy based on two-dimensional cross-sectional slices from XCT tomograms was implemented, substantially increasing the number of training samples while preserving relevant structural information.

Two prediction strategies were evaluated: a displacement-derived approach from our previous D^2^IM and a direct strain prediction model D^2^IM-Strain. The results showed that directly learning strain fields from the input tomograms improves prediction performance, particularly for strain magnitudes below the yield threshold of bone in compression (i.e. 10000με). Specifically, the direct strain model significantly reduced false positive classifications of high-strain regions, achieving a 75% reduction compared with the displacement-derived approach. This indicates a stronger ability to distinguish low-strain regions while maintaining reliable identification of high-strain areas. These findings suggest that directly predicting strain fields from imaging data can improve the reliability of strain estimation while avoiding the additional processing steps required in displacement-derived methods. Future work will explore multi-modal physics-informed neural network architectures [41–43] to improve physical plausibility and extrapolation beyond training data distributions, combining the flexibility of data-driven learning with the interpretability of mechanics-based models. Through these developments, the D^2^IM framework has the potential to advance the mechanical characterization of bone and other hierarchical materials, providing rapid, accurate, and noise-resistant strain predictions that enhance our understanding of structure-function relationships across scales.

## Data availability statement

Code for preparing dataset, training D^2^ IM model and visualising/analysing results has been hosted on GitHub: https://github.com/M34Impact/D2IM-Strain

The dataset used for this study can be found on Figshare: https://doi.org/10.6084/m9.figshare.25404220.v1

## Author statement

**Jon Valijonov, Peter Soar, James Le Houx and Gianluca Tozzi:** Conceptualisation of the methodology, data analysis, and drafting, editing, reviewing and finalising the paper. **Jon Valijonov and Peter Soar:** Digital volume correlation analysis, software/code development and execution, modelling, data collection and visualisation.

## Acknowledgements

The authors acknowledge support from the School of Engineering and the School of Computing and Mathematical Sciences within the Faculty of Engineering and Science at the University of Greenwich. This research received no external funding.

